# Can genomics and meteorology predict outbreaks of legionellosis in urban settings?

**DOI:** 10.1101/2022.09.26.509626

**Authors:** Verlaine J Timms, Eby Sim, Keenan Pey, Vitali Sintchenko

## Abstract

*Legionella pneumophila* is ubiquitous and sporadically infects humans causing Legionnaires disease (LD). Globally, reported cases of LD has risen four-fold from 2000-2014. In 2016, Sydney, Australia was the epicentre of an outbreak caused by *L. pneumophila* serogroup 1 (Lpsg1). Whole genome sequencing was instrumental in identifying the causal clone which was found in multiple locations across the city. This study examined the epidemiology of Lpsg1 in an urban environment, assessed typing schemes to classify resident clones and investigated the association between local climate variables and LD outbreaks. Of 223 local Lpsg1 isolates, we identified dominant clones with one clone isolated from patients in high frequency during outbreak investigations. The cgMLST scheme was the most reliable in identifying this Lpsg1 clone. While an increase in humidity and rainfall was found to coincide with a rise in LD cases, the incidence of the major *L. pneumophila* outbreak clone did not link to weather phenomena. These findings demonstrated the role of high resolution typing and weather context assessment in determining source attribution for LD outbreaks in urban settings, particularly when clinical isolates remain scarce.

**Importance:** We investigated the genomic and meteorological influences of infections caused by *Legionella pneumophila* in Sydney, Australia. Our study contributes to a knowledge gap of factors that drive outbreaks of legionellosis compared to sporadic infections in urban settings. In such cases, clinical isolates can be rare and other data is then relied upon to inform decision making around control measures. We found that cgMLST typing offered a robust and scalable approach for high-resolution investigation of Lpsg1 outbreaks. The genomic landscape of Lpsg1 in Sydney was dominated by a single clone which was responsible for multiple clusters of community cases over four decades. While legionellosis incidence peaked in Autumn, this was not linked to the dominant outbreak clone. The synthesis of meteorological data with Lpsg1 genomics can be a part of the risk assessment for legionellosis in urban settings and is relevant for other densely populated areas around the world.

## Introduction

In 2016 Sydney was the epicentre of several outbreaks of legionellosis that spanned three distinct geographical regions with the same clone found to be responsible for all three outbreaks (*1*). The reasons for this apparent connection remain unclear.

*Legionella pneumophila* is ubiquitous in the environment and (*2*) is responsible for cases of legionellosis, which encompass Legionnaire’s disease (LD), a severe pneumonia and Pontiac fever, a self-limiting, influenza like illness in humans. Cases of legionellosis peak in autumn or summer but consistently in times of higher rainfall and humidity (*3,4,5*), with a lag time of 1-2 weeks (*4,6*). Legionellosis is diagnosed more often in males over 50 years of age, especially smokers or those with underlying health conditions (*4,5*). The incidence of legionellosis has substantially increased for all age groups in the US since 1990 (*4,7*) and in Hong Kong a >4-fold increase has occurred since 2005 (*5*). The incidence rate of legionellosis has more than tripled in the US between 2000-2011 (*4*) and a similar trend has been noted in Europe since 2004 (*8*). The cause of increasing incidence is currently not explained but may be related to the growing numbers of susceptible individuals at high risk for LD due to age and comorbidities, improved diagnosis due to the availability of the urinary antigen test, increased use of syndromic PCR panels and better reporting to national registries (*4*).

Despite a LD case-fatality rate of 30% (*7*) public health and laboratory capacity to identify LD outbreaks remains limited. Obstacles include, but are not limited to, obtaining adequate clinical samples for testing, isolating *L. pneumophila* from respiratory samples and appropriate environmental sampling and testing. Further, once an isolate has been recovered and identified, the resolution of serogroup typing of *L. pneumophila* is limited, as among 16 serogroups of *L. pneumophila*, serogroup 1 (Lpsg1) is responsible for the majority (80-84%) of human disease (*7*). A sequence based typing (SBT) scheme improves the resolution of *L. pneumophila* typing (*9*). However, not all Lpsg1 strains can be readily typed with this method when whole genome sequencing (WGS) is employed (*9,10*). With WGS offering the ultimate resolution for molecular epidemiology a detailed understanding of the genomic variability of Lpsg1 in the environment is essential for accurate interpretation of genetic relatedness.

This study examined the epidemiology of Lpsg1 clones in an urban environment and assessed the ability of typing schemes to classify resident clones. We employed SBT, Bayesian analysis of population structure (BAPS) led hierarchical clustering and core genome multi-locus sequence typing (cgMLST) to identify and characterise the major disease-causing clones in Sydney. We also investigated the effect of environmental factors, such as temperature and rainfall on LD outbreaks. To achieve this, we sequenced all available isolates since 1982 to determine the genomic landscape of Lpsg1 in Sydney as an example of a contemporary urban environment.

## Methods

### Bacterial isolates, serogrouping and whole-genome sequencing

*Legionella pneumophila* serogroup 1 isolates referred to the Centre for Infectious Diseases and Microbiology Laboratory Services, NSW Health Pathology between 1982 and 2019, collected as sporadic cases and outbreak associated investigations were included. Two reference strains of Lp1, ATCC33152 and ATCC33215, were added to represent global strains (Supplementary Table). Isolates were grown and serotyped as described previously using standard methods (*1*). Genomic DNA was extracted and sequenced on the Illumina platform as previously described (*1*).

### Genome assembly and phylogenetic analysis

The raw Fastq files were assessed for accurate basecall quality (Phred score over 20) and presence of adaptors using FastQC (https://www.bioinformatics.babraham.ac.uk/projects/fastqc/). Kraken (1.0) (*11*) was used to confirm the reads were from *Legionella* sp. Sequencing reads were assembled with Spades (3.9.0) (*12*) and assembly quality was assessed using Quast (version 5.0.2) (*13*). Assemblies were annotated with Prokka (version 1.13.3).

Pangenome assessment was performed using the Roary pipeline (3.6.1) (*14*) including core genome alignment using Multiple Alignment using Fast Fourier Transform (MAFFT) (7.300) (*15*). This core genome alignment was used for building a maximum likelihood (ML) phylogeny with 1000 bootstrap replicates and the GTR+F+R5 model as determined by ModelFinder (16) in IQ-TREE (1.6.7) (*17*). The core genome alignment from Roary was used as the input into Bayesian analysis of population structure (BAPS) (*18*) using iterative clustering to a depth of 10 levels and a prespecified maximum of 20 clusters.

### Genome typing

Sequence based typing (SBT) was performed on the assemblies with seven established loci (*9*) using Legsta (0.5.1) (https://github.com/tseemann/legsta) which contained a database of 2,793 SBT sequence types (ST). Genomes not designated a ST were assigned a probable ST if they possessed an allelic profile identical to six loci but had variation (inclusive of missing data/novel calls) in the *mompS* locus. If the variation was not in *mompS* and had an allelic number assigned to the variable loci, these genomes were designated as a unique ST. If any genomes had more than two loci with either missing data or novel calls not localised in *mompS*, they were designated as untypeable. The core genome Multi Locus Sequence Typing (cgMLST) schema of *Legionella pneumophila* hosted on Ridom (https://www.cgmlst.org/ncs/schema/1025099/; accessed 01/12/2020) was adapted for use in chewBACCA version 2.5.5. Each genome was assigned an 18-mer relative neighbourhood address (CGRNA), consisting of six tiers in the format of “NNN-NNN-NNN-NNN-NNN-NNN” whereby each “NNN” represented an individual tier. Tiers were clustered based on the number of allelic differences between genomes. Genomes with more than 1000 allelic differences were separated and assigned a three-digit numeric code, forming the first tier. All genomes within a tier were further sequentially sub-clustered based on thresholds of 150, 15, 5, 2 and 0 allelic differences with each decreasing threshold constituting a different tier and assigned a three-digit numeric code. This segregation allowed for six modular levels with CGRNA-6 incorporating all six tiers with sequential decrease in levels reflective of the number of tiers used. Results of the cgMLST was visualised as a minimum spanning tree on Grapetree (*19*) using the MSTreeV2 algorithm.

### Meteorologic data and case modelling

Daily weather observations were provided by the Australian Bureau of Meteorology with data spanning 1990-2019 at the Observatory Hill station located in Sydney. Monthly legionellosis notifications for New South Wales were obtained from the National Notifiable Disease Surveillance System (NNDSS) and aggregated alongside monthly means of 14-day-lagged daily weather observations at Observatory Hill. Visualisations of mean weather statistics and case counts were generated to observe previously reported seasonal effects of LD case counts. A quasi-Poisson linear regression model was input with NNDSS case counts and mean monthly weather observations. The model was iteratively trained using a leave-one-out approach to determine the predictive value over each year. The model was optimised and variables selected according to the QAIC parameter available in R package MuMIn (https://rdrr.io/cran/MuMIn/).

### Genome typing and weather observations

The cgMLST barcode of all 223 isolates were paired to dates of specimen collection for meteorologic analyses. As these dates do not reliably represent date of symptom onset, both 7- and 14-day lag periods were tested to account for the time between symptom onset and microbiological testing. Weather observations for lagged dates were compared across cgMLST barcode types at varying resolutions and overlaid on weather observation figures.

### Data Availability

The genomic data have been deposited in the NCBI Sequence Read Archive (SRA) (http://www.ncbi.nlm.nih.gov/Traces/sra/) under Bioproject ID PRJNA767747 with the Lpsg1 genomes from 2016 under study accession number SRP117289.

## Results

A total of 223 Lpsg1 isolates from Sydney were sequenced, including 160 clinical and 63 environmental isolates with 40 identified as epidemiologically associated with outbreak investigations. Sequences were assembled with the average genome size in the range of 3.2-3.8 Mb (Supplementary Table), a core genome (>95%) of 1231 genes and an accessory genome (<95%) of 9750 genes. Further statistics on assembly quality including N50 and contig number are provided in the Supplementary Table.

BAPS clustering was able to assign every isolate to a cluster (Figure 1 and Supplementary Table) and three dominant clusters were identified. The largest cluster was Cluster 4 88/233 (39%), while 63/223 (28%) were assigned to Cluster 6 and 42/223 (19%) were assigned to Cluster 1. To investigate the context of the three major BAPS clusters seen in Sydney, a maximum likelihood phylogeny was inferred from an alignment of 1231 core genes. The resultant phylogenetic tree was overlaid with year collected, BAPS cluster and if the isolate was collected as part of an outbreak investigation (Figure 1). Dominant BAPS clusters co-occurred across the time of the study while isolates recorded as part of an outbreak occurred in BAPS Cluster 4.

**Figure 1:**
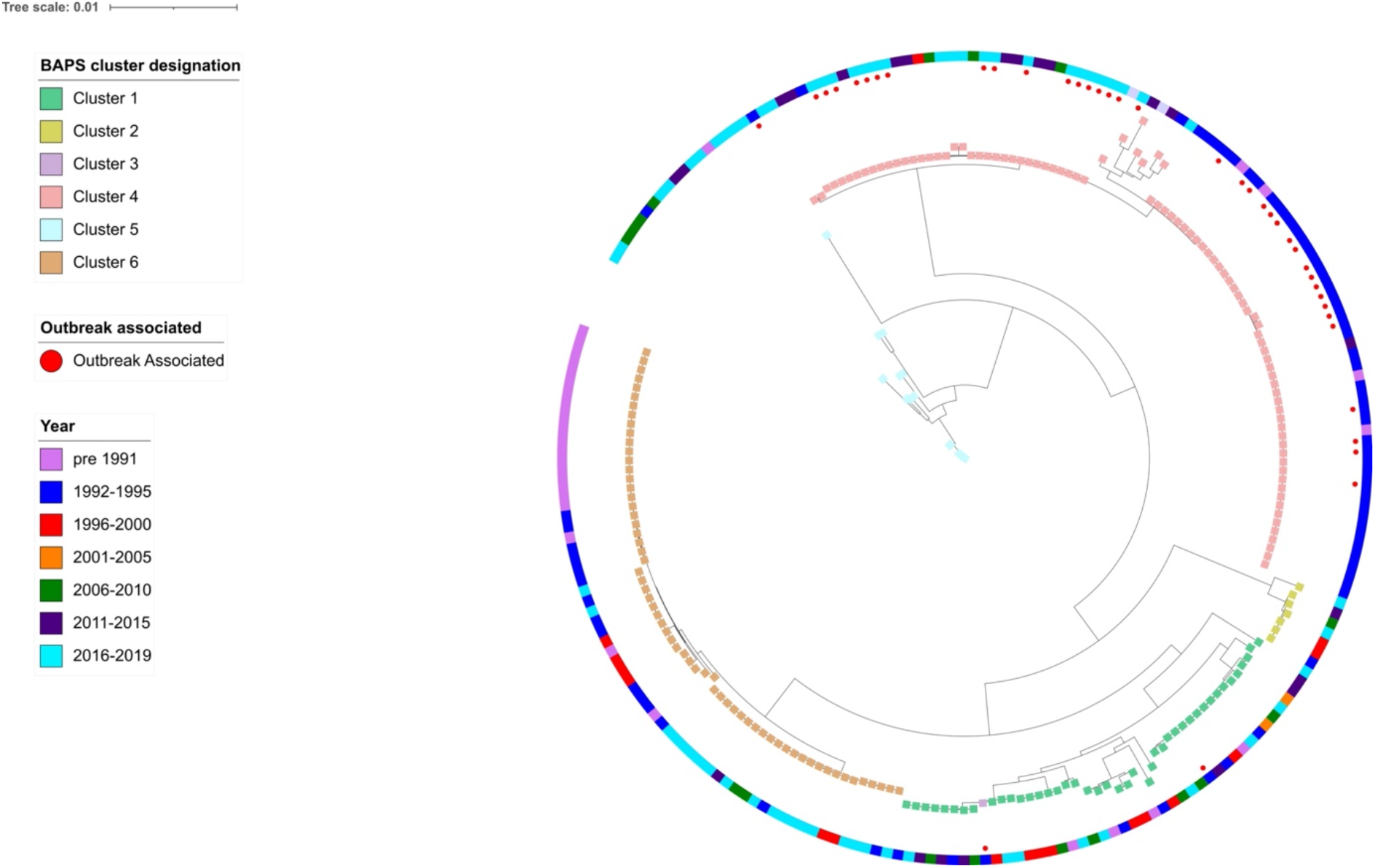
Core genome phylogeny of 223 Lpsg1 genomes based on 1231 core genes. Coloured squares indicate BAPS cluster designation. Red dots indicate whether the isolate was collected as part of an outbreak investigation. Colour bar on the perimeter indicates year collected in 5-year increments according to the legend. A higher density of isolates recorded as part of an outbreak occurs in BAPS cluster 4. Visualisation produced in iTOL (https://itol.embl.de/).

### Lpsg1 genome typing

Using the SBT method, 160/223 genomes (71.75%) were successfully typed without further interrogation. Genomes belonging to ST37 formed the majority (43/223; 18.45%) of the dataset followed by ST1 (33/223; 14.80%) and ST211 (31/223; 13.90%). The rest of the typeable Lpsg1 genomes were spread over 18 different STs. Of the 63 genomes that required further interrogation, two were untypeable with each having either novel or missing data over two loci, 34 genomes could be assigned to a probable ST according to definitions outlined in the methods, while the remaining 27 genomes were designated as novel STs, spread over 13 different ST types.

Each *L. pneumophila* genome was individually assigned a relative neighbourhood address after cgMLST which resulted in 161 unique cgMLST relative neighbourhood addresses (CGRNA; Level 6; Supplementary Table) based on number of allelic differences between genomes. Visualisation on a minimum spanning tree provided a snapshot of the finer genomic diversity of Lpsg1 in Sydney, a scale that could be overlooked from just SBT and a high-level scan of BAPS designation (Figure 2). At a cut-off threshold of 1,000 allelic differences (CGRNA-1), all 223 Lpsg1 were segregated unevenly into 12 different CGRNA-1 neighbourhoods with 9 (90/233; 38.63%), 8 (63/233; 27.04%) and 7 (42/233; 18.03%) forming the top-3 CGRNA-1 neighbourhoods. CGRNA-1 neighbourhood 7 was observed to be the most diverse with some genomes showing 474-922 allelic difference between them. A similar observation was also noted in CGRNA-1 neighbourhoods 3 and 12 where a single isolate in each neighbourhood possessed more than 900 allelic differences relative to its closest neighbour.

**Figure 2:**
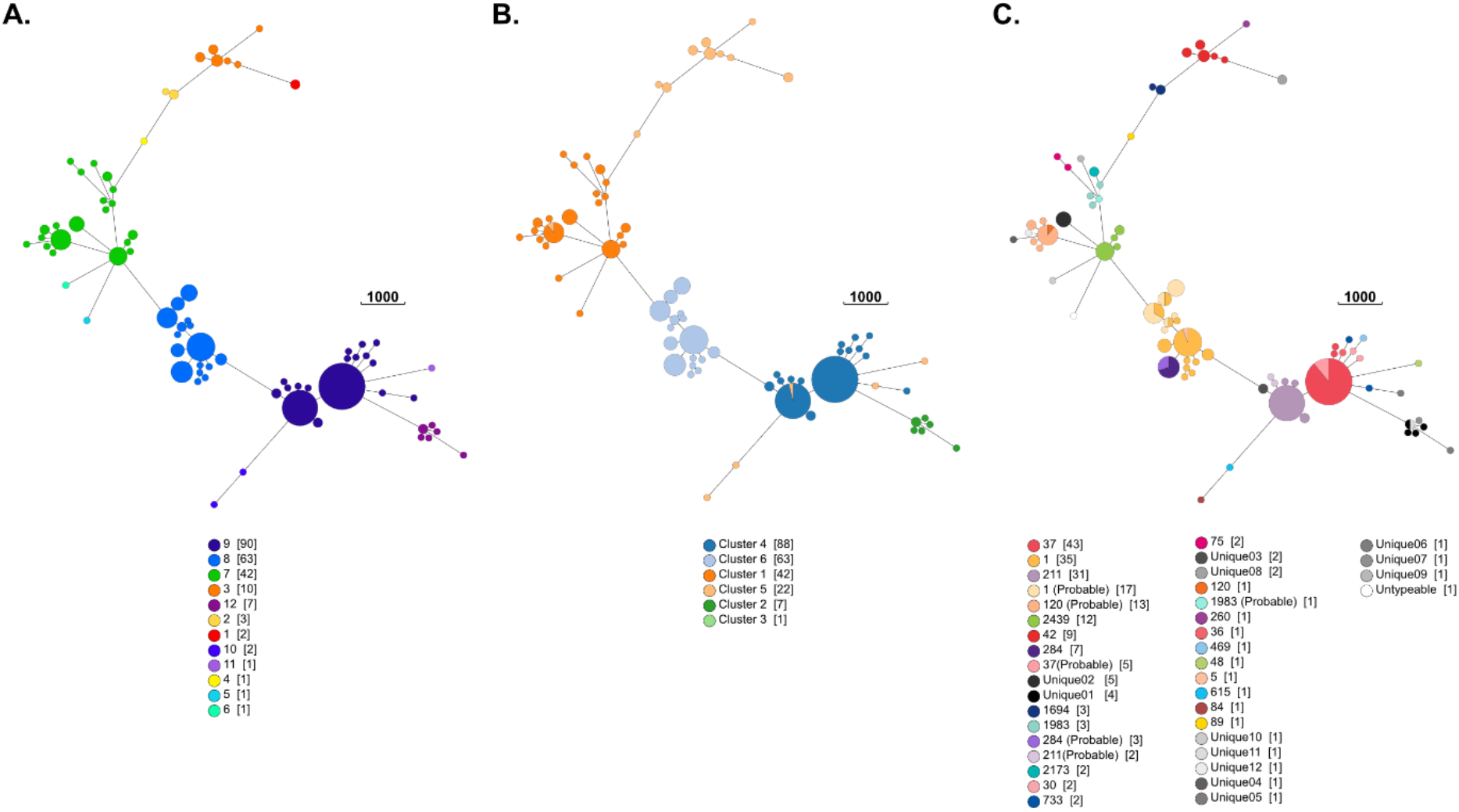
Minimum spanning tree of the 223 Lpsg1 genomes based on 1,521 loci with genomes with 5 allelic difference (CGRNA-4) collapsed and overlaid with (A) CGRNA-1 (B) BAPs cluster (C) SBT sequence types. Each node represents either a single or a collection of genomes, colour coded with pertinent information based on the different overlaid information. Allelic differences are scaled according to the scale bar. Visualisation produced with Grapetree (19).

The utilisation of cgMLST also allowed for complementation of data as clustering results of the adapted cgMLST schema was generally in concordance with both BAPS and SBT (Figure 2B & C) designations. With regards to BAPS and cgMLST concordance, genomes belonging to BAPS cluster 1,2,4 and 6 appeared stratified within their associated CGRNA-4 clusters (Figure 2B). We are unable to comment upon the sole genome designated as BAPS cluster 3 but the only exception to the aforementioned stratification were some genomes (n=3) that belonged to BAPS cluster 5 which clustered amongst other genomes belonging to CGRNA-4 neighbourhoods stratified by BAPS 1 (n=1) and BAPS 4 (n=2).

Similar to BAPS, SBT designations were also stratified within their respective CGRNA-1 neighbourhood with no STs found in multiple neighbourhoods (Figure 2C). This also included genomes designated as “probable” by SBT, which similarly clustered together with their associated counterparts. Of the 12 unique ST designated in this study, only genomes that belonged to ST Unique11 were clustered together with genomes belonging to a different ST (ST37) within similar CGRNA-4. This observation was not entirely unexpected as both ST Unique11 and ST37 differed by a single locus with genomes that belonged to the former having a novel *neuA* allele, not represented in the Legsta database. This was also observed in another genome which was designated as untypeable by SBT. In addition to possessing a novel *neuA*, untypeable by Legsta, this particular genome possessed a novel allele at *mompS* and was thus classified as an untypeable. Our results (Figure 2) indicated that both cgMLST and SBT were generally concordant with each other but complete concordance was not observed between cgMLST and BAPs due to the aforementioned BAPS 5 designated genomes.

### Meteorologic data and case modelling

The linear model revealed key components of the relationship between LD and weather. The monthly linear model suggested significant associations between LD cases and mean relative humidity at 9am and 3pm (p<0.005), and average temperature at 9am and 3pm (p<0.05) (Figure 3). Rainfall, cloud coverage, sunshine hours, and previous month statistics did not show any correlation with LD case counts.

**Figure 3.**
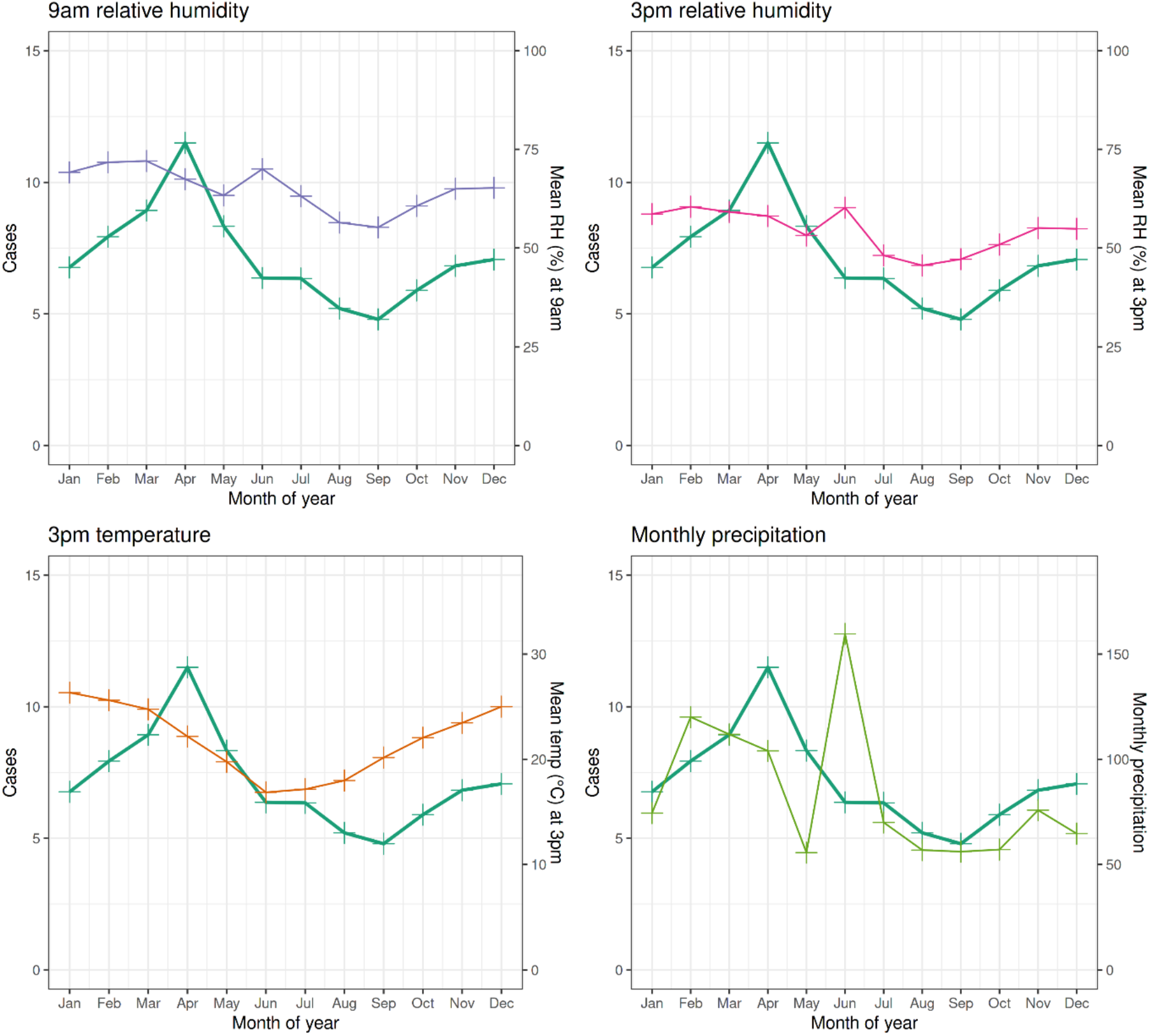
Mean monthly weather observations versus reported LD case counts across NSW over the calendar year. Data ranges from years 1990 to 2019. LD case count means are represented statically across all four subfigures in green. Top left: relative humidity at 9am, blue. Top right: relative humidity at 3pm, pink. Bottom left: air temperature at 3pm, orange. Bottom right: monthly precipitation, green. Results visualised with ggplot2 (20).

### Genome typing and weather observations

When the CGRNA-5 scheme was applied, limiting the number of displayed categories to seven, outbreak types 009-001-002-001-004 and 009-001-002-001-002 were found to occur together temporally in line with observations of BAPS cluster and year of collection (Figures 1 and S1).Predominant outbreak CGRNA-5 009-001-002-001-004 and 009-001-002-001-002 appeared over multiple varying weather modes, representing a range of +/-10 degrees and +/- 25% relative humidity at 9am. Additionally, sporadic cgMLST isolates persist in all but extreme weather conditions, an observation consistent with the collection dates of sporadic cases across the calendar year (Figure 4).

**Figure 4.**
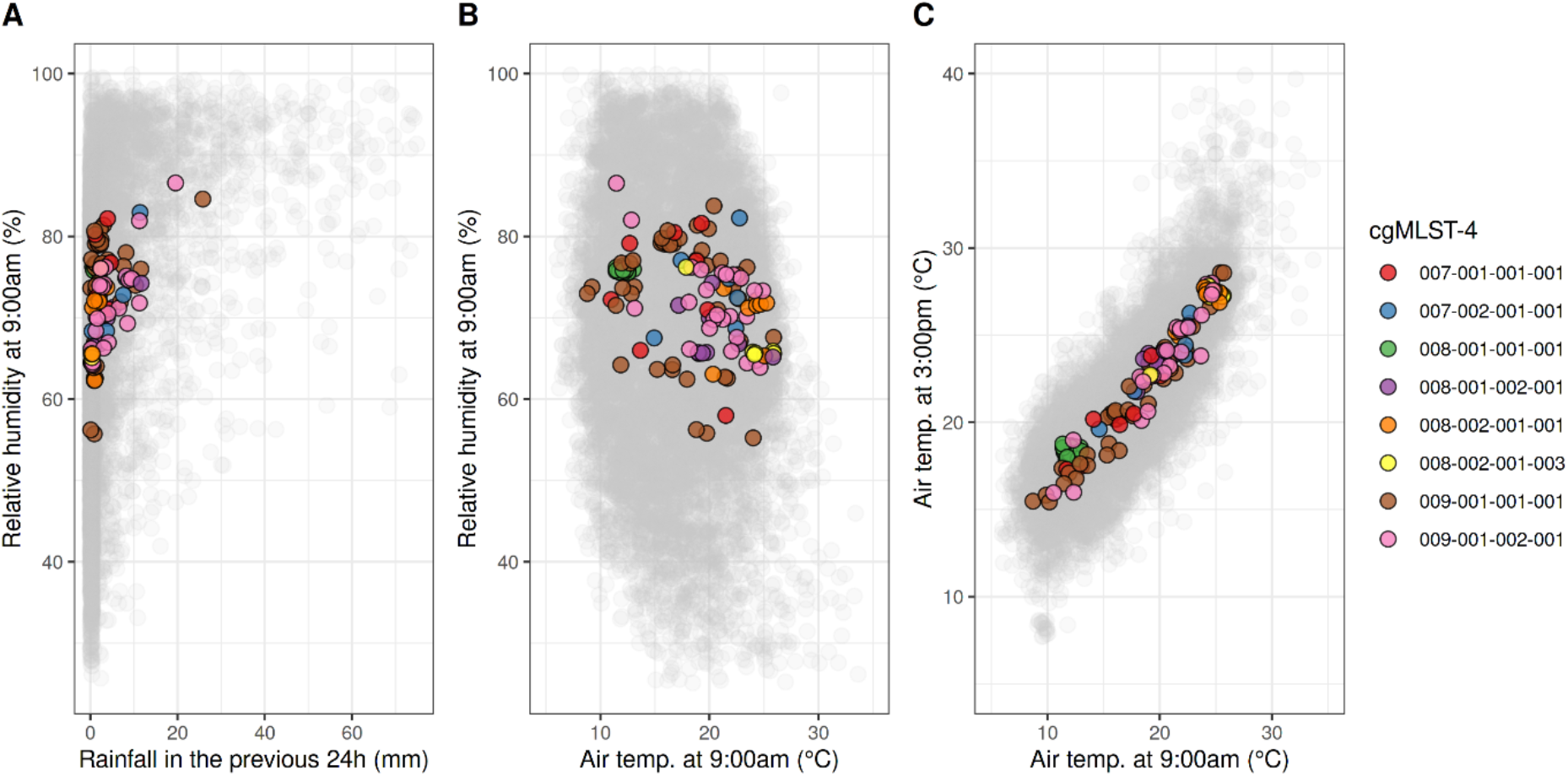
14-day lagged weather observations versus 14-day lagged moving average weather at time of isolate collection by cgMLST. Individual cgMLSTs are indicated by coloured circles. (**A**) Relative humidity at 9:00am versus total rainfall in the previous 24 hours. (**B**) Relative humidity versus air temperature at 9:00am. (**C**) Air temperature at 9:00am versus air temperature at 3:00pm. All circles are randomly jittered for clarity of isolate group sizes. Grey circles represent dates on which no isolates in this study were collected. Results visualised with ggplot2 (20).

## Discussion

This study interrogated 223 isolates of Lpsg1 representing evolving urban populations of *L. pneumophila* in Sydney over a period of nearly 40 years. The motivation was to investigate the genomics of local strains responsible for LD with the view to improve molecular surveillance and early detection of outbreaks. One dominant clone of Lpsg1 appeared to be responsible for three large outbreaks while sporadic cases were usually explained by other regularly occurring clones. Two typing methods were used to distinguish clones and a cgMLST scheme was devised that capitalised on the high resolution of WGS data. Given weather conditions have a demonstrated effect on the incidence of *L. pneumophila* infections in other cities around the world, local weather conditions were investigated and found to indeed impact LD incidence but not appropriately forecast the occurrence of major outbreaks.

BAPS clustering provided a population snapshot of the 233 Lpsg1, assigning the genomes into six different population clusters based on the core gene content. In addition to BAPS clustering, we also performed *in silico* typing utilising both SBT and a publicly available cgMLST scheme. Both schemes were each capable of adding further resolution compared to BAPS clustering, at least at the population clustering level. SBT has been the mainstay subtyping methodology for well over a decade (*9,21*). However, the increasing ubiquity of second-generation sequencing platforms has led to the ease of generating WGS data and as such, a cgMLST scheme was recently developed for *L. pneumophila* typing (*10*). As a typing tool, cgMLST has been shown to be a scalable approach for high resolution analysis (*22*) and therefore was compared to the SBT typing scheme.

A common feature of all three methods employed in this study was the utilisation of draft assemblies as input, which are then assessed with either a panel of preselected loci (cgMLST and SBT) or a panel derived from the current dataset (core genome alignment followed by BAPS). In addition, the software(s) utilised for each typing scheme can omit nuances of possible non-assembly due to either coverage issues over the loci, a repeated allele (e.g., *mompS*), or G+C bias. This highlighted the need for some manual curation of the outputs over these automated processes. While non-assembly of regions of the genome can impact upon the clustering results, both cgMLST (and the derived CGRNA) and BAPS clustering employed in study are able to provide some mitigation, at the expense of resolution, due to the tiered clustering approach and core genome generation respectively.

Notwithstanding the possible impacts of sequencing artefacts, our findings still documented a significant degree of concordance between BAPS, SBT and cgMLST typing schemes in the classification of the Lpsg1 genomes. Complete concordance between BAPS and cgMLST however, was not observed. While both methods were fundamentally similar, each utilising a set of loci to segregate genomes, there are also differences within their implementation. The adapted cgMLST scheme has a fixed number of loci and did not require annotation of the genomic assemblies while in this BAPS analysis, an alignment of assembled and annotated genomes was used as input. The cgMLST scheme is based on a select group of closed genomes some of which may not be reflective of Australian clones while BAPS clustering was derived from local draft genomes. It was therefore possible that differences in the respective schemas led to the observation of non-concordance in six legionella genomes. On the other hand, genomes belonging to the same/and or closely similar ST were all stratified in their associated CGRNA neighbourhood. This observation echoed previous reports that SBT could be a reliable proxy for genetic lineage, even within the WGS era (*23*). However, a previously reported limitation of SBT when WGS data was used as input is due to some isolates harbouring two copies of *mompS*. Similarly, the majority of our genomes that were not typed showed either a failure of detection of the *mompS* locus via *in silico* PCR or variation in this locus (*24*). While we tentatively assigned these genomes into probable ST based on cut-offs declared in this study, the application of cgMLST further indicated that these assigned probable STs could be in fact the correct STs due to the higher discriminative resolution of cgMLST. Despite some minor discrepancies between these three methods, the general concordance between them did echo previous suggestions that cgMLST can be a “gold standard” for *L. pneumophilia* typing, if access to WGS data is assured (*10*).

Meteorologic data in the context of environmentally mediated infections like LD has been shown to be an important area of cross-disciplinary research (*25*). Given the observed phenomena of LD attributed to weather conditions in other countries, we sought to provide a glimpse of this attribution in a local context and to provide an empirical basis for heuristics that describe LD appearing at Easter time. Linear modelling failed to capture a link between weather conditions and high-count months, suggesting either incorrect model inputs and parameters or, perhaps more likely, additional factors at play in the development of LD. The sparse nature of the underlying legionellosis case count data may serve as a contributing factor to this failure, despite the use of “quasi-Poisson” linear regression modelling suiting a rarely occurring event such as LD. However, frequent parity was seen between model predicted counts and months of low, sporadic cases. This suggests the utility of meteorologic data in predicting case counts lies in the ability to anticipate sporadic cases but not outbreaks.

The “genometeoromics” approach to the multi-disciplinary combination of meteorologic and genomic data revealed no association between virulent outbreak types and the weather. We set out to determine the existence of temperature or humidity “cut-offs” beyond which outbreak-associated strains are found in order to develop a heuristic for increased public health sampling. The genomic and weather comparison in Figure 4 demonstrated a lack of weather pattern driving particular Lpsg1 genotypes to cause human disease. The wide dispersion of cases from many weather points suggests a lack of connection between genotype and specific weather patterns, instead indicating non-meteorologic factors are primarily at play in the establishment of LD outbreaks. This coincides with the linear model’s inability to capture weather information relevant to upticks in case counts and outbreaks. Together both results form a narrative that the incidence of *L. pneumophila* outbreak versus sporadic strains is not inherently linked to weather phenomena.

The use of NNDSS case counts for these analyses carries with it an important caveat - case counts are reported by state, but weather observations are limited to a centre located within Sydney. The state of New South Wales is estimated as 801,150 square kilometres in area, with two thirds of the population residing in the greater Sydney region. This presents, assuming an even distribution of LD cases across the population, that weather observations do not accurately reflect conditions for one third of the population of LD cases.

While this study rests on a considerable number of Lpsg1 strains isolated over nearly 40 years, an important limitation to note is that clinical Lpsg1 isolates represent a small percentage of those patients suffering from LD and an even smaller percentage of the Lpsg1 genetic pool that exists in the environment. Developments in direct metagenomic sequencing have the potential to resolve problems isolating clinical strains so that future population studies will provide a more complete snapshot of the strains that cause human disease. Environmental Lpsg1 are sampled extensively during outbreaks and rarely in non-outbreak periods. As such the potential flux of Lpsg1 strains in the environment is unknown although some evidence of wide variability in environmental populations exists (*26*). Studies that sample and investigate Lpsg1 strains from environmental sources across seasons will help to close this gap in knowledge.

In conclusion, the cgMLST typing offers a robust and scalable approach for high-resolution detection and investigations of Lpsg1 outbreaks in urban settings. The genomic landscape of Lpsg1 in Sydney has been dominated by a single clone, which has been responsible for multiple clusters of community cases over four decades. While legionellosis incidence was found to peak in Autumn this was not linked to the dominant outbreak clone. The synthesis of meteorological data, such as rainfall and outdoor temperature, with Lpsg1 genomics can be a part of the risk assessment for legionellosis in urban settings and can be applied in other densely populated areas around the world.

## Acknowledgements

This research was supported by the Sydney Informatics Hub and the University of Sydney’s high performance computing service, Artemis. The authors wish to thank the Pathogen Genomics Team and the Enteric Reference Laboratory at the Centre for Infectious Diseases and Microbiology Laboratory Services – New South Wales Health Pathology for performing genome sequencing. Funding from Population Health and Health Services Research Support Program, NSW Health, Australia.

## Figures

**Supplementary Table: Collection data and typing of 223 genomes of *L. pneumophila***. Tab 1 Sample information including source, outbreak, year isolated and BAPS cluster. Tab 2 Assembly statistics including length, N50, GC% and number of contigs. Tab 3 SBT typing results included alleles. Tab 4 CGRNA designations.

## References

1. Timms VJ, Rockett R, Bachmann NL, Martinez E, Wang Q, Chen SC-A, et al. Genome sequencing links persistent outbreak of legionellosis in Sydney (New South Wales, Australia) to an emerging clone of Legionella pneumophila sequence type 211. Appl Environ Microbiol. 2018;84(5).

2. Ditommaso S, Giacomuzzi M, Memoli G, Garlasco J, Zotti CM. Persistence of legionella in routinely disinfected heater-cooler units and heater units assessed by propidium monoazide QPCR. Pathogens. 2020;9(11):1–14.

3. Sakamoto R. Legionnaire ‘ s disease, weather and climate. Bull World Heal Organ. 2015;93(January):435–6.

4. Hicks LA, L G, Ge N, Hampton LM. Legionellosis - United States, 2000-2009. MMWR Morbitity Mortal Wkly Rep. 2011;60:1083.

5. Leung Y. Epidemiology of Legionnaires’ disease, Hong Kong, China, 2005-2015. Emerg Infect Dis. 2020;26(8):1695.

6. Fisman DN, Lim S, Wellenius GA, Johnson C, Britz P, Gaskins M, et al. It’s not the heat, it’s the humidity: Wet weather increases legionellosis risk in the greater Philadelphia metropolitan area. J Infect Dis. 2005;192(12):2066–73.

7. Mercante JW, Morrison SS, Desai HP, Raphael BH, Winchell JM. Genomic Analysis Reveals Novel Diversity among the 1976 Philadelphia Legionnaires’ Disease Outbreak Isolates and Additional ST36 Strains. PLoS One. 2016;11(9):e0164074.

8. Beauté J. Legionnaires’ disease in Europe, 2011 to 2015. Eurosurveillance. 2017;22(27):1–8.

9. Gaia V, Fry NK, Afshar B, Lück PC, Meugnier H, Etienne J, et al. Consensus sequence-based scheme for epidemiological typing of clinical and environmental isolates of Legionella pneumophila. J Clin Microbiol. 2005 May [cited 2016 Oct 14];43(5):2047–52.

10. Moran-Gilad J, Prior K, Yakunin E, Harrison TG, Underwood A, Lazarovitch T, et al Design and application of a core genome multilocus sequence typing scheme for investigation of Legionnaires’ disease incidents. Eurosurveillance. 2015;20(28):21186.

11. Wood DE, Salzberg SL. Kraken: Ultrafast metagenomic sequence classification using exact alignments. Genome Biol. 2014;15(3).

12. Bankevich A, Nurk S, Antipov D, Gurevich AA, Dvorkin M, Kulikov AS, et al. SPAdes: A New Genome Assembly Algorithm and Its Applications to Single-Cell Sequencing. J Comput Biol. 2012;19(5):455–77.

13. Gurevich A, Saveliev V, Vyahhi N, Tesler G. QUAST: Quality assessment tool for genome assemblies. Bioinformatics. 2013;29(8):1072–5.

14. Page AJ, Cummins CA, Hunt M, Wong VK, Reuter S, Holden MTG, et al. Roary: rapid large-scale prokaryote pan genome analysis. Bioinformatics. 2016;31(22):3691–3.

15. Katoh K, Misawa K, Kuma K-I, Miyata T. MAFFT: a novel method for rapid multiple sequence alignment based on fast Fourier transform. Nucleic Acids Res. 2002;30(14):3059–66.

16. Kalyaanamoorthy S, Minh BQ, Wong TKF, von Haeseler A, Jermiin LS. ModelFinder: fast model selection for accurate phylogenetic estimates. Nat Methods. 2017;14(6):587–9.

17. Nguyen LT, Schmidt HA, Von Haeseler A, Minh BQ. IQ-TREE: A fast and effective stochastic algorithm for estimating maximum-likelihood phylogenies. Mol Biol Evol. 2015;32(1):268–74.

18. Cheng L, Connor TR, Sirén J, Aanensen DM, Corander J. Hierarchical and spatially explicit clustering of DNA sequences with BAPS software. Mol Biol Evol. 2013/02/13. 2013 May;30(5):1224–8.

19. Zhou Z, Alikhan NF, Sergeant MJ, Luhmann N, Vaz C, Francisco AP, et al. Grapetree: Visualization of core genomic relationships among 100,000 bacterial pathogens. Genome Res. 2018;28(9):1395–404.

20. Wickham H. ggplot2: Elegant Graphics for Data Analysis. Springer-Verlag New York; 2016. Available from: https://ggplot2.tidyverse.org

21. Ratzow S, Gaia V, Helbig JH, Fry NK, Lück PC. Addition of neuA, the gene encoding N-acylneuraminate cytidylyl transferase, increases the discriminatory ability of the consensus sequence-based scheme for typing Legionella pneumophila serogroup 1 strains. J Clin Microbiol. 2007;45(6):1965–8.

22. Pearce ME, Alikhan N-F, Dallman TJ, Zhou Z, Grant K, Maiden MCJ. Comparative analysis of core genome MLST and SNP typing within a European Salmonella serovar Enteritidis outbreak. Int J Food Microbiol. 2018; 274:1–11.

23. Underwood AP, Jones G, Mentasti M, Fry NK, Harrison TG. Comparison of the Legionella pneumophila population structure as determined by sequence-based typing and whole genome sequencing. BMC Microbiol. 2013;13(1):302.

24. Gordon M, Yakunin E, Valinsky L, Chalifa-Caspi V, Moran-Gilad J. A bioinformatics tool for ensuring the backwards compatibility of Legionella pneumophila typing in the genomic era. Clin Microbiol Infect. 2017 May;23(5):306–10.

25. Walker JT. The influence of climate change on waterborne disease and Legionella: a review. Perspect Public Health. 2018;138(5):282–6.

26. Wüthrich D, Gautsch S, Spieler-Denz R, Dubuis O, Gaia V, Moran-Gilad J, et al. Air-conditioner cooling towers as complex reservoirs and continuous source of Legionella pneumophila infection evidenced by a genomic analysis study in 2017, Switzerland. Eurosurveillance. 2019;24(4):1–7.

